# Replication Kinetics, Pathogenicity and Virus-induced Cellular Responses of Cattle-origin Influenza A(H5N1) Isolates from Texas, United States

**DOI:** 10.1101/2024.10.29.620905

**Authors:** Ahmed Mostafa, Ramya S. Barre, Anna Allué-Guardia, Ruby A. Escobedo, Vinay Shivanna, Hussin Rothan, Esteban M. Castro, Yao Ma, Anastasija Cupic, Nathaniel Jackson, Mahmoud Bayoumi, Jordi B Torrelles, Chengjin Ye, Adolfo García-Sastre, Luis Martinez-Sobrido

## Abstract

The host range of HPAIV H5N1 was recently expanded to include ruminants, particularly dairy cattle in the United States. Shortly after, human H5N1 infection was reported in a dairy worker in Texas following exposure to infected cattle. Herein, we rescued the cattle-origin influenza A/bovine/Texas/24-029328-02/2024(H5N1, rHPbTX) and A/Texas/37/2024(H5N1, rHPhTX) viruses, identified in dairy cattle and human, respectively, and their low pathogenic forms, rLPbTX and rLPhTX, with monobasic HA cleavage sites. Intriguingly, rHPhTX replicated more efficiently than rHPbTX in mammalian and avian cells. Still, variations in the PA and NA proteins didn’t affect their antiviral susceptibility to PA and NA inhibitors. Compared to rHPbTX and rLPbTX, the rHPhTX and rLPhTX exhibited higher pathogenicity and efficient replication in infected C57BL/6J mice. The lungs of rHPhTX-infected mice produced higher inflammatory cytokines/chemokines than rHPbTX-infected mice. Our results highlight potential risk of HPAIV H5N1 virus adaptation in human and/or dairy cattle during the current multistate/multispecies outbreak.

## Introduction

In late 2020, a novel reassortant of highly pathogenic avian influenza A virus (HPAIV) H5N1 clade 2.3.4.4b was detected in Europe and became the most predominant reassortant/variant in Europe in poultry and wild birds [1]. One year later, HPAIV H5N1 was carried by migratory birds across the Atlantic Ocean to North America, including the United States (US) [1,2]. Since then, HPAIV H5N1 clade 2.3.4.4b outbreaks in avian populations (wild and domestic) and infections of humans, carnivores, herbivores and marine mammals have been frequently reported.

Influenza viruses, including influenza A (IAV), B (IBV), C (ICV), and D (IDV), are identified as members of the family *Orthomyxoviridae* with different host ranges for each type [3]. IAV genome is composed of eight negative-sense and single-stranded RNA segments, including the three polymerase-encoding segments and their accessory proteins (PB2, PB1, PB1-F2, PA and PA-X), hemagglutinin (HA), neuraminidase (NA), nucleoprotein (NP), matrix proteins-encoding segment (M1 and M2), and nonstructural proteins-encoding segment (NS1 and NEP) [3].

Since 2011, positive cattle for IDV have been detected in multiple geographic areas across the world [4]. Until March 2024, cattle were only proposed as the primary reservoir of IDV [4]. In late March 2024, this classification was amended after the recent transmission of HPAIV H5N1 of clade 2.3.4.4b (genotype B3.13) in dairy cattle in the US via milking equipments [5–7]. Subsequently, HPAIV H5N1 was transmitted to cats, raccoons, opossums, mice and birds on affected dairy farms, probably via contaminated splashed milk. Recent studies have shown that the mammary glands are the primary sites that support efficient viral replication [6,8] and HPAIV H5N1 titers in milk could rapidly peak to 10^8^ TCID_50_/mL following intramammary inoculation [6].

From March to October 2024, 10 human infections with HPAIV H5N1 have been reported to be associated with exposures to infected dairy cattle with conjunctivitis but mild respiratory symptoms [9,10]. One scenario that may explain the ocular and mild respiratory symptoms is that some farm workers get infected after a direct splashing of contaminated milk from infected cattle to their eyes [10].

The pathogenicity of IAV is a complex and multigenic trait that is affected by several viral and host factors [3]. Intensive circulation of HPAIV H5N1 in new animal reservoirs like cattle can result in the emergence of genetic traits with sufficient capability to cross the animal-to-human barrier(s) and improve its fitness (e.g. PB2 with 627K substitution) [3]. These genetic changes could be acquired in the new (human) host or the intermediate animal host (cattle). The impact of these changes in the virus phenotype and its zoonotic potential for adaptation and transmission among humans needs to be investigated [3].

In late March 2024, an adult dairy farm worker in an affected dairy farm in Texas had an onset of redness and discomfort in the right eye [11]. The case was further reported as the first human case of cattle-flu-related HPAIV H5N1, and the infecting virus was named A/Texas/37/2024(H5N1). Compared to the closely related co-circulating cattle-flu isolates (e.g. cattle-origin influenza A/bovine/Texas/24-029328-02/2024(H5N1), the human isolate was accompanied by distinct nonsynonymous mutations in PB2, PB1, PA, NA and NS segments of unknown characteristics in mammals. Herein, we constructed, using reverse genetic systems, recombinant A/Texas/37/2024(H5N1; rHPhTX) and A/bovine/Texas/24-029328-02/2024(H5N1; rHPbTX) to investigate the impact of their amino acid (aa) variations on viral replication kinetics, susceptibility to FDA-approved influenza antivirals, virus-induced host responses, and pathogenicity in C57BL/6J mice. We further generated low pathogenic forms of A/Texas/37/2024(H5N1; rLPhTX) and A/bovine/Texas/24-029328-02/2024(H5N1; rLPbTX) with a monobasic cleavage site in the viral HA and investigated the possible synergistic impact of amino acids (aa) variations in the different PB2, PB1, PA, NA and NS segments, and the HA multibasic cleavage site, *in vitro* and *in vivo*. The rHPhTX showed higher replication kinetics than the bovine rHPbTX in different mammal and avian cell types. Notably, the rHPhTX replicated more efficiently in the lungs, nasal turbinate, and brains and showed higher pathogenicity in C57BL/6J-infected mice than rHPbTX. This increase in viral pathogenicity of rHPhTX compared to rHPbTX can be, at least in part, attributed to the higher cytokine storm observed in the lungs of infected C57BL/6J mice. Moreover, the rLPhTX and rLPbTX were attenuated compared to their rHPhTX and rHPbTX counterparts, with rLPhTX replicating more efficiently and causing higher pathogenicity than the rLPbTX. Importantly, rHPhTX and rHPbTX showed similar susceptibility to PA (Baloxavir) and neuraminidase (Oseltamivir and Zanamivir) inhibitors, demonstrating the feasibility of using FDA-approved antivirals for the treatment of these new HPAIV A(H5N1) infections of both cattle and human origin.

## Materials and Methods

### Biosafety

All the *in vitro* and *in vivo* experiments with highly pathogenic and low pathogenic avian influenza H5N1 viruses were conducted in appropriate biosafety level (BSL) 3 and animal BSL3 (ABSL3) laboratories, respectively, at Texas Biomedical Research Institute (Texas Biomed). Experiments were approved by the Texas Biomed Institutional Biosafety (IBC) and Animal Care and Use (IACUC) committees.

### Cells and Viruses

Madin-Darby canine kidney (MDCK), human embryonic kidney (293T), Madin-Darby bovine kidney (MDBK), human lung adenocarcinoma epithelial (A549), Crandell-Rees Feline Kidney (CRFK), and chicken fibroblast (DF-1) cells were maintained in Dulbecco’s modified Eagle medium (DMEM) (Invitrogen, Carlsbad, CA, USA) supplemented with 10% fetal bovine serum (FBS) and 1% PSG (penicillin, 100 U/mL; streptomycin 100 μg/mL; l-glutamine, 2 mM) at 37°C in a humidified 5% CO_2_ incubator. MDCK cells were kindly provided by Daniel Perez at the Animal Health Research Center, Center for Vaccines and Immunology, Department of Population Health, Poultry Diagnostic and Research Center. The highly pathogenic cattle-origin human influenza A/Texas/37/2024 (H5N1, HPbTX) virus was provided by Todd Davis and Han Di at the Virology Surveillance and Diagnosis Branch, Influenza Division, The Centers for Disease Control and Prevention (CDC).

### Cloning and generation of recombinant viruses

The viral segments of the human influenza virus A/Texas/37/2024 (H5N1, HPhTX) were synthesized in the genetic background of pUC57 (GenScript, USA). The synthesized sequences were designed according to the published sequences for the clinical HPhTX isolate with GenBank accession numbers PP577940-47, respectively. The synthesized viral segments were further subcloned into linearized pHW2000 to construct pHW-hTX_PB2, pHW-hTX_PB1, pHW-hTX_PA, pHW-hTX_HA, pHW-hTX_NP, pHW-hTX_NA, pHW-hTX_M and pHW-hTX_NS [12,13].

To generate the pHW-bTX_PB2, pHW-bTX_PB1, pHW-bTX_PA, pHW-bTX_NA and pHW-bTX_NS plasmids expressing the viral segments of recombinant highly pathogenic influenza A/bovine/Texas/24-029328-02/2024(H5N1, HPbTX), the corresponding pHW-hTX_PB2, pHW-hTX_PB1, pHW-hTX_PA, pHW-hTX_NA and pHW-hTX_NS plasmids were subjected to site-directed mutagenesis using specific primers (**Table 1**). The sequences of viral HA, NP, and M segments of HPhTX and HPbTX are identical.

**Table 1.**
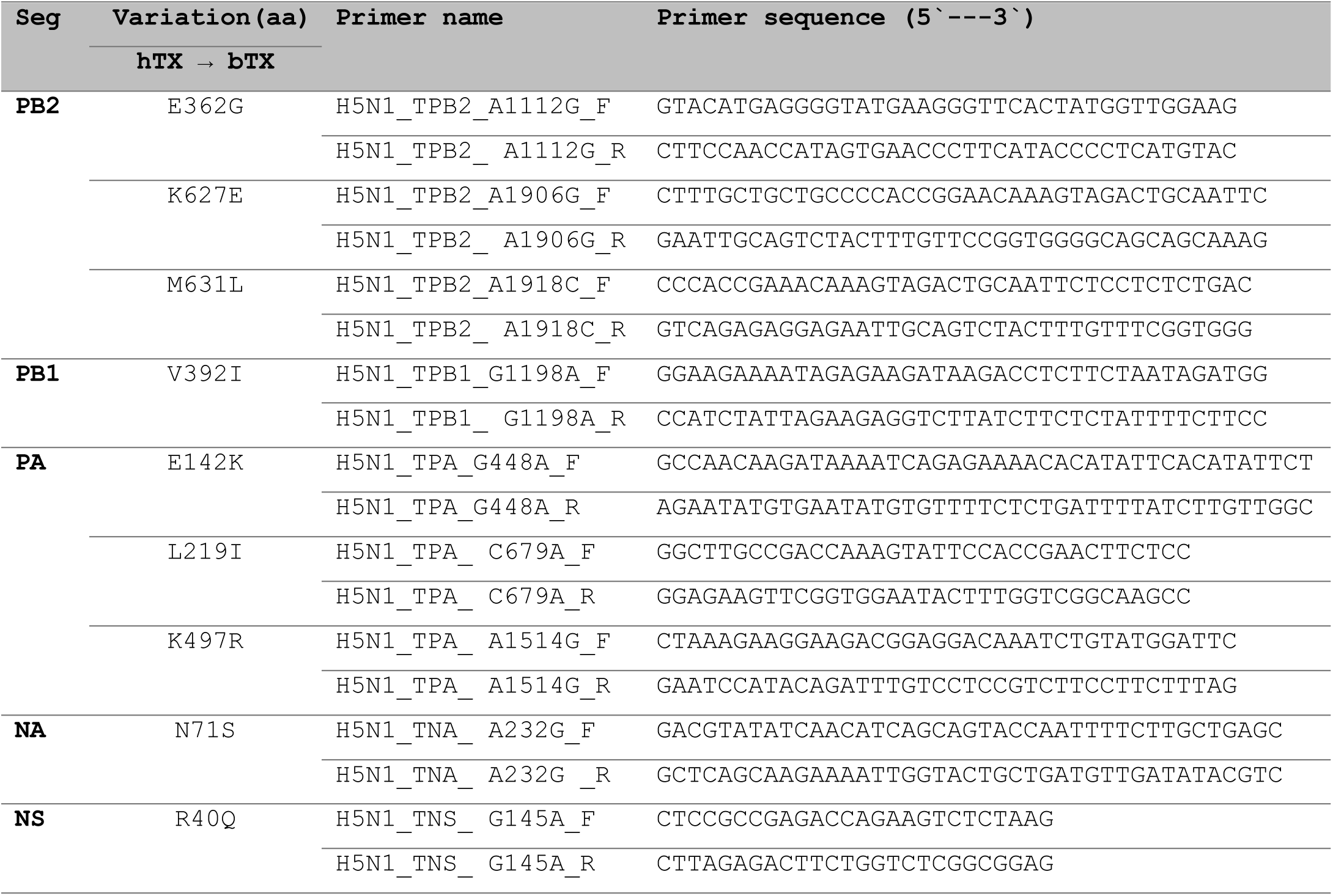
Primers used to generate pHW plasmids to rescue rHPbTX.

The multibasic cleavage site (PLREKRRKR/GLF) in the viral HA was converted to a monobasic cleavage site (PQIETR/GLF) by amplifying the HA1 and HA2 separately using outer universal primers [14] and internal primers (Bm_LPH5HA_2344R1: 5’-TATTCGTCTCTTCTCGTTTCTATTTGAGGACTATTTCTGA-3’ & Bm_LPH5HA_2344F2: 5’-TATTCGTCTCGGAGAGGTCTGTTTGGGGCGATAGCAG-3’) as previously described [15]. Briefly, 2 μl (5 ng/μl) of pHW-hTX_HA were mixed with 5 μl 10X *Pfu* reaction buffer, 1 μl 10mM dNTP mixture (0.2 mM each), 1.5 μl of 50 mM MgCl2 (1.5 mM), 2 μl of each Primer (40 pmoles) and 1 μl of *PfuTurbo* DNA Polymerase (2.5 U/μl) (Invitrogen, USA). The total volume was adjusted to 50 μl using nuclease-free water (Ambion, USA), followed by pre-denaturation at 95°C for 2 min, PCR amplification (35 cycles: 95°C/30 sec for denaturation, 56°C/30 sec for annealing and 72°C/3 min for extension), then final extension at 72°C/10 min. The PCR products were purified using “Wizard® SV Gel and PCR Clean-Up System” (Promega, USA). The obtained RT-PCR fragments were next digested with BsmBI-v2 (NEB, USA) and cloned into pHW2000 as previously described to generate pHW-LPhTX_HA.

To generate rHPhTX, rLPhTX, rHPbTX, and rLPbTX, the relevant set of the eight ambisense pHW2000 plasmids (1 μg each) was co-transfected into a co-culture of 293T/MDCK-II cells (ratio 3:1) using Lipofectamine™ 3000 Transfection Reagent (ThermoFisher Scientific, USA) according to manufacturer instructions. Following transfection, the media was replaced with 1 mL of “Opti-MEM” containing penicillin/streptomycin and 0.2% bovine serum albumin (BSA) and incubated in a humidified 5% CO_2_ incubator for 12 h. Afterwards, an additional 1 mL of “Opti-MEM” containing 1% P/S/G and 0.2% BSA was added to each well. For rLPhTX and rLPbTX, an additional 1 mL of “Opti-MEM” containing 1% P/S/G and 0.2% BSA was supplemented with 2 μg/mL TPCK-treated trypsin (Sigma–Aldrich, USA). At 48–72 h post-addition of TPCK-treated trypsin, the cell culture supernatants were harvested and centrifuged at 2,500 rpm for 5 min at 4°C. A portion of the collected cell culture supernatant was then used to infect MDCK monolayers in T-75 cell culture flask in the presence of infection media (DMEM containing 1% P/S/G and 0.2% BSA with (rLPhTX and rLPbTX) or without (rHPhTX and rHPbTX) 1 μg/mL TPCK-treated trypsin. The recombinant viruses (rHPhTX, rLPhTX, rHPbTX, and rLPbTX) were then aliquoted and stored at −80°C until use. All viruses were confirmed by whole genome sequencing of the viral stocks using the next-generation sequencing platform, MinION (Oxford Nanopore Technologies). Viral RNA was extracted using the QiAamp Viral RNA Mini Kit (Qiagen). Sample libraries were prepared using the Native Barcoding Kit 24 V14 (SQK-NBD114.24, Oxford Nanopore Technologies) and ran on R10.4.1. Flow Cells (FLO-MIN114, Oxford Nanopore Technologies) per the manufacturer’s instructions. All viruses were corroborated/compared with the published clinical HPhTX isolate (GenBank accession numbers: PP577940-47).

### Plaque Assay and Immunostaining

MDCK cells were cultured at the density of 10^6^ cells per well in 6-well plates and kept overnight at 37°C in a humidified 5% CO_2_ incubator. The next day, confluent MDCK monolayers were infected with serial viral dilutions for 1 h at 37°C. Following viral adsorption, cell monolayers were overlaid with post-infection media containing agar and incubated at 37°C in a humidified 5% CO_2_ incubator. At 24-72 hours post-infection (hpi), cells were fixed overnight with 10% neutral buffered formalin solution. For staining and visualization using crystal violet, 1 mL of 1% crystal violet solution was added to each well for 5 min at room temperature (RT) and then rinsed with tap water. For immunostaining, cells were permeabilized with 0.5% (vol/vol) Triton X-100 in PBS for 15 min at RT and immunostained using an influenza virus anti-NP mouse monoclonal antibody (MAb) HT103 (1:100) and the Vectastain ABC kit (Vector Laboratories), following the manufacturers’ instruction. After immunostaining, plates were scanned and photographed using a ChemiDoc MP Imaging System.

### Viral replication kinetics

Monolayers of A549, CRFK, MDBK, and DF-1 cells were cultured in 6-well plates (10^6^ cells per well) for 24 h, infected in triplicate with indicated viruses at a multiplicity of infection (MOI) of 0.001 and then kept at 37°C in a humidified 5% CO_2_ incubator to allow viral adsorption for 1 h. Following viral adsorption, the infected cell monolayers were washed three times with 1X PBS to remove residual non-adsorbed viral particles. The plates were then incubated at 37°C in a humidified 5% CO_2_ incubator. The cell culture supernatants were collected at 12, 24, 48, and 72 hpi. Viral titers in collected samples were determined as previously described using plaque assay and immunostaining [16].

### Antiviral susceptibility

MDCK cells were seeded into 96-well plates at a density of 2×10^4^ cells per well and incubated at 37°C for 24 h. After removing the growth media, cells were washed once with PBS and 100 particle-forming units (PFU) of the virus were added per well. Cells were incubated for 1 h at 37°C in a humidified 5% CO_2_ incubator to allow viral adsorption. After incubation, the virus inocula were removed, and 100 µl of infection media containing varying concentrations of each compound were added. Treated and control infected MDCK monolayers were then incubated for 48 h. Following incubation, cells were fixed with 10% neutral buffered formalin solution and stained with 1% crystal violet solution. Once dried, 200 µL of 100% methanol was added to each well, and absorbance was measured at 560 nm. The EC_50_ values were calculated to identify the concentration required to elicit 50% of the maximal response. To determine the EC_50_, concentration-response data were analyzed using GraphPad Prism 9.5.1 software. A non-linear regression model, log(agonist) vs. normalized response—variable slope, was applied to fit the data.

### Mice experiments

Female C57BL/6J mice (n=13/group) were anesthetized intraperitoneally (i.p.) by a cocktail of Ketamine (100 mg/mL) and Xylazine (20 mg/mL). Anesthetized mice were infected intranasally (i.n.) with the indicated viral doses in a total volume of 50 μl PBS. At days 2 and 4 post-infection (dpi), 4 mice from each group were euthanized to collect lung, nasal turbinate, brain, heart, kidney, spleen, and mammary gland tissues. The half organ was fixed in 10% neutral buffered formalin solution for histopathology and immunohistochemistry (IHC) analyses, and the other half was homogenized in 1 mL of PBS using a Precellys tissue homogenizer (Bertin Instruments) for viral titration. Tissue homogenates were centrifuged at 10,000 *xg* for 5 min and the supernatants were used to determine viral load and investigate the presence of secreted chemokines/cytokines. The remaining 5 animals were monitored for 14 days for disease progression, body weight changes, and mortality/survival rates. Mice that experienced a weight loss exceeding 25% of their original weight were humanely euthanized.

### Multiplex cytokine assay

Cytokines and chemokines (IFN-α, IFN-β, IFN-γ, IL-6, TNF-α, MCP-1 or CXCL2, IP-10 or CXCL-10, and RANTES or CCL5) were measured using a custom 8-plex panel mouse ProcartaPlex assay (ThermoFisher Scientific, cat. number: PPX-11-MXMFZCY, lot number: 417770-000), following the manufacturer’s instructions. Lung homogenate samples were centrifuged at 10,000 *xg* for 5 min prior to performing the assay to remove debris and diluted 1:2 in Universal Assay Buffer. The assay was performed in the ABSL-3 laboratory, and samples were decontaminated by overnight incubation in 1% formaldehyde solution before readout on a Luminex 100/200 System running on xPONENT v4.3.309.1 with the following parameters: gate 7,500-25,000, 50 μl of sample volume, 50 events per bead, sample timeout 60 s, Standard PMT. Acquired data were analyzed using xPONENT v4.3.309.1.

### Histopathology and immunohistochemistry (IHC)

Mice lungs fixed in 10% neutral buffered formalin solution were embedded in paraffin, sectioned and stained with hematoxylin and eosin (H&E). Briefly, the processing of formalin-fixed tissues occurred in a Tissue Tek VIP tissue processor by dehydration through a series of graded alcohols, clearing through one change of 50:50 absolute alcohol/xylene mixture and two changes of xylene, and concluded with the infiltration of tissues with paraffin wax using ParaPro™ XLT infiltration and embedding media (StatLab, USA). The paraffin-embedded blocks were cut using a Microm HM325 rotary microtome at a thickness of 4 microns and mounted onto microscope slides using a flotation water bath set between 46° and 48°C. Slides were loaded onto the Varistain Gemini automated slide stainer for H&E staining and dried in heating stations before the staining program proceeded. The deparaffinization of tissue sections occurred through three changes of xylene, two changes of absolute alcohol, and two changes of 95% alcohol and rinsed in distilled water. Hematoxylin (StatLab, USA) staining commenced, followed by the removal of excess stain using High Def solution (StatLab, USA) and bluing of the hematoxylin stain using Reserve Bluing Reagent (StatLab, USA). Tissue sections were then stained with Eosin (StatLab, USA), followed by dehydration in three changes of alcohol, a 50:50 alcohol/xylene mixture, and cleared in three changes of xylene before coverslipping. Upon completion of all staining, tissue sections were evaluated using light microscopy by a board-certified veterinary pathologist.

Tissue sections for IHC were cut at 4 microns thick, mounted onto positively charged slides and allowed to air dry overnight. Slides were then loaded onto the Discovery Ultra IHC/ISH automatic stainer for the detection of the avian influenza virus. Deparaffinization occurred using Discovery Wash (Roche, USA). Cell conditioning occurred using Discovery CC1 (Roche, USA) at 95°C for 64 minutes (min). The blocking of endogenous peroxidase occurred using Discovery Inhibitor (Roche, USA) for 8 min. Slides were then incubated with an influenza A NP rabbit polyclonal antibody (Invitrogen, USA) at a concentration of 1:1500 for 1 h at RT and detected using anti-rabbit HQ (Roche, USA) for 8 min, followed by Anti-HQ HRP (Roche, USA) for 8 min at 36° C. Influenza A NP was then visualized using ChromoMAP DAB (Roche, USA). The slides were counterstained using Hematoxylin (Roche, USA) followed by Bluing Reagent (Roche, USA).

### Quantification and statistical analysis

Graphing and statistical analyses were performed using GraphPad Prism 8 (GraphPad Software, San Diego, California, USA; www.graphpad.com). The unpaired, two-tailed Student’s t-test was used for two-group comparisons for each time-point and reported as p < 0.05; p < 0.005; p < 0.0005. Multiple comparisons among groups and/or time points were analyzed using analysis of variance (ANOVA) with Tukey’s post hoc test. For cytokine and chemokine analyses, a two-way ANOVA with Holm-Sidak’s multiple comparisons test was used to compare groups within each time point. Statistical significance was reported as *p < 0.05; **p < 0.005; ***p < 0.0005; ****p < 0.00005. The “ns” indicates non-significance.

## Results

### Sequence analyses of HPhTX and HPbTX and generation of recombinant viruses

The HPhTX and HPbTX were isolated in Texas in April 2024 from human and dairy cattle, respectively. Compared to HPbTX, the HPhTX has several amino acid (aa) variations in the PB2 (G362E, E627K and L631M), PB1 (I392V), PA (K142E, I219L and R497K), PA-X (K142E), NA (S71N), and NS1 (Q40R) proteins (**Fig 1a**). Interestingly, HPhTX contains the PB2 viral mutation E627K while the HPbTX has PB2 viral mutation M361L present in ≥99% of dairy cattle isolates (**Fig 1a**). Both aa substitutions in HPhTX (E627K) and HPbTX (M361L) were previously reported to be associated with viral adaptation to mammalian hosts, including humans [17,18]. Compared to other published viral protein sequences of human H5N1 strains that were detected in the US since April 2024, these nine aa substitutions in HPhTX were distinct (**Fig 1b**) and three of them (PA_219L, PA_497K and NA_71N) were occasionally identified in the US human H5N1 isolates since April 2024 (**Supplemental data**). Conversely, the HA, NP, M1, M2, and NS2 viral proteins were identical in both HPhTX and HPbTX.

**Figure 1.**
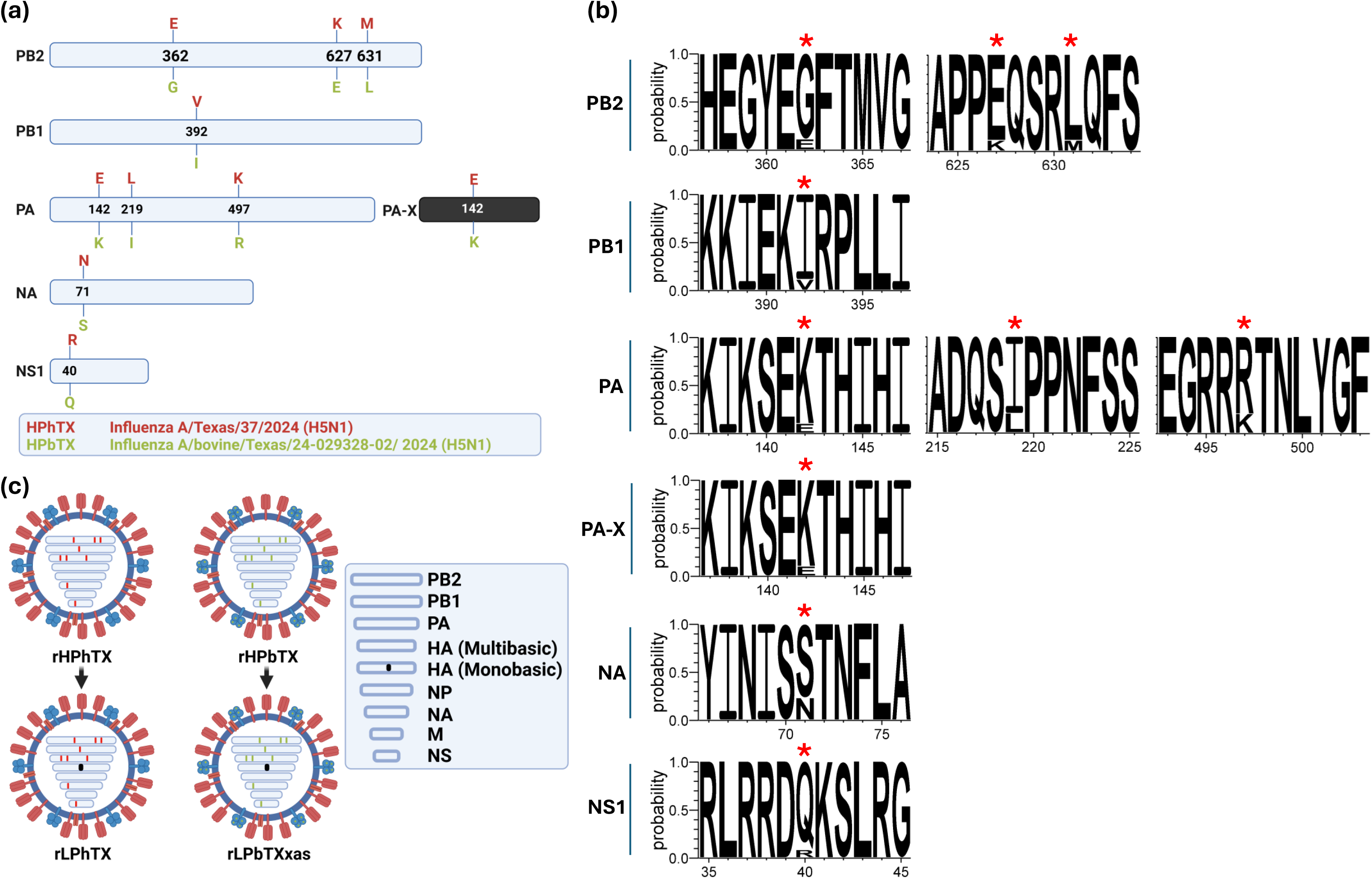
Sequence analysis of rHPhTX and rHPbTX. (**a**) Schematic presentation of the amino acid (aa) variations between rHPhTX (red, top) and rHPbTX (green, bottom) in PB2, PB1, PA, PAX, NA, NS1 proteins (top to bottom). (**b**) Prevalence rate of the distinct aa residues in HPhTX among recently reported human isolates in the US since April 2024. The graphic was created via a Web-based WebLogo application (http://weblogo.threeplusone.com/create.cgi) [35]. Red asterisks indicate which aa are prevalent among the human H5N1 isolates in the US since the cattle H5N1 outbreak in March 2024. (**c**) Schematic representation of the rHPhTX (top left), rHPbTX (top right), rLPhTX (bottom left), and rLPbTX (bottom right) viruses used in this study. The figure has been created with BioRender.com.

The reverse genetics (RG) approach enables the rescue of recombinant influenza A viruses using cloned cDNA plasmids [13,15,19]. We employed this approach to generate the recombinant HPhTX (rHPhTX) that shares 100% protein homology with the published clinical isolate of HPhTX. Using site-directed mutagenesis (**Table 1**), we next constructed the RG system of the closely related HPbTX (rHPbTX) to compare back-to-back with the recombinant human isolate (rHPhTX). Next, recombinant low pathogenic forms of HPhTX and HPbTX were rescued, namely rLPhTX and rLPbTX, by changing the multibasic cleavage site of the viral HA to a monobasic cleavage site (**Fig 1c**). To confirm that the rHPhTX was phenotypically similar to the clinical natural isolate HPhTX, both viruses were compared for their plaque phenotype and replication kinetics in mammalian and avian cell lines. Indeed, both viruses showed comparable plaque phenotype (**Fig 2a**) and growth kinetics in A549 (human), MDBK (bovine), CRFK (feline), and DF-1 (avian) cells (**Fig 2b**).

**Figure 2.**
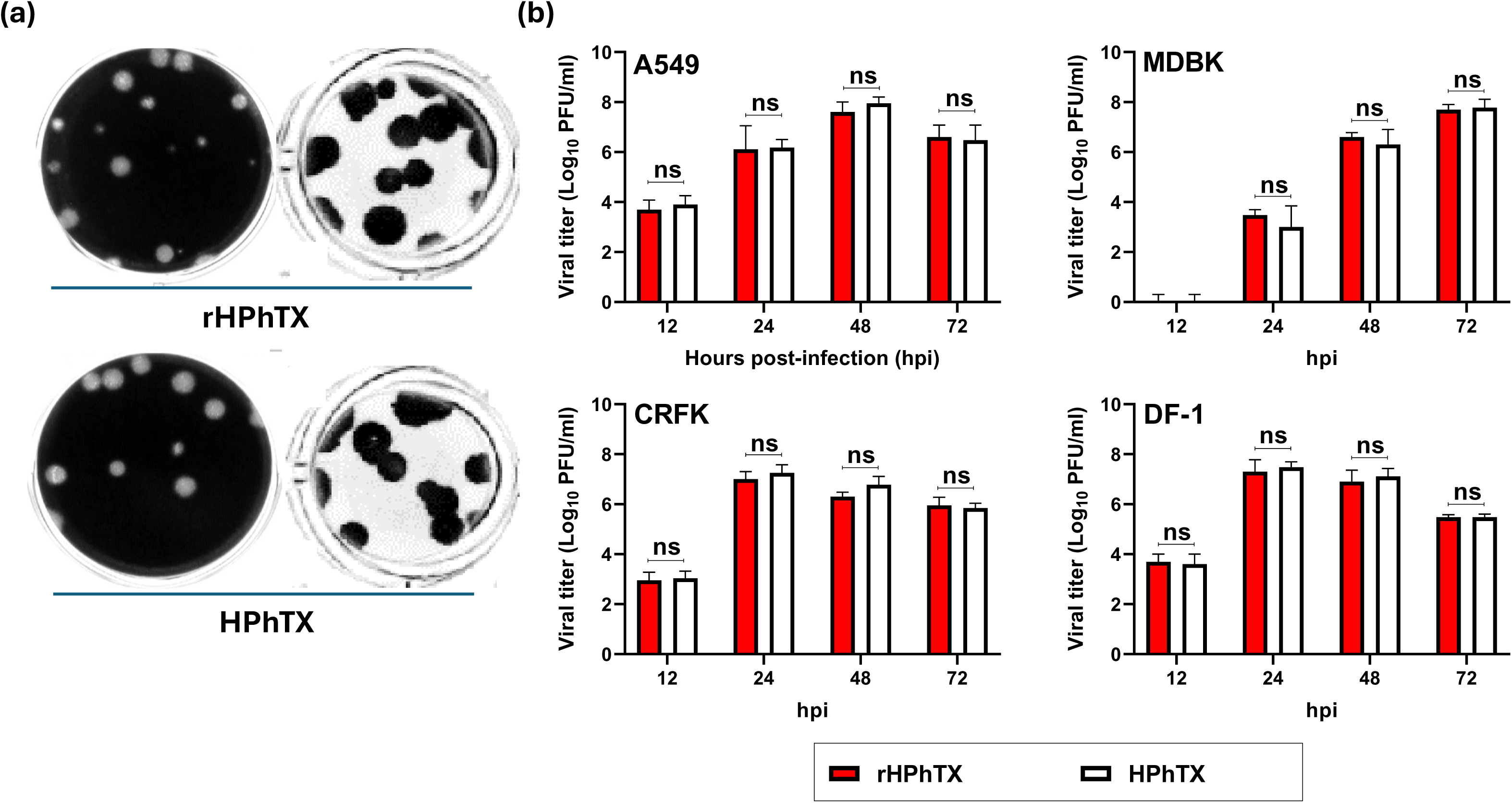
Phenotypic characterization of the rHPhTX and the natural HPhTX isolate. (**a**) Plaque morphology of rHPhTX (top) and the natural HPhTX isolate (bottom) using crystal violet (left) and immunostaining (right). (**b**) Replication kinetics of rHPhTX (red) and HPhTX natural isolate (white) in A549, MDBK, CRFK, and DF-1 cells. Error bars reflect the standard deviation (SD) of three independent replicates. Statistical analysis was performed using repeated measures ANOVA, followed by Bonferroni *post hoc* test. The “ns” indicates a non-significant difference.

Next, we confirmed the low pathogenicity of the generated rLPhTX and rLPbTX by comparing them to rHPhTX and rHPbTX in the presence and absence of 1 μg/mL TPCK-treated trypsin using plaque assay (**Fig 3a**). Unlike the rHPhTX and rHPbTX that can efficiently form viral plaques in the absence of TPCK-treated trypsin, the rLPhTX and rLPbTX could form viral plaques only in the presence of TPCK-treated trypsin. We also investigated the preferential replication of rHPhTX and rHPbTX in mammalian and avian cell lines (**Fig 3b**). Compared to the rHPbTX, the rHPhTX replicated to significantly higher titers in A549, CRFK, MDBK, MDCK, and MDBK cells, especially at early time points post infection (12-24 hpi) (**Fig 3b**). This suggests that the aa differences between rHPhTX and rHPbTX impact the replication of both viruses in mammalian and avian cells, with better replication of rHPhTX, although differences in replication appeared to be smaller in avian DF-1 cells (**Fig 3b**).

**Figure 3.**
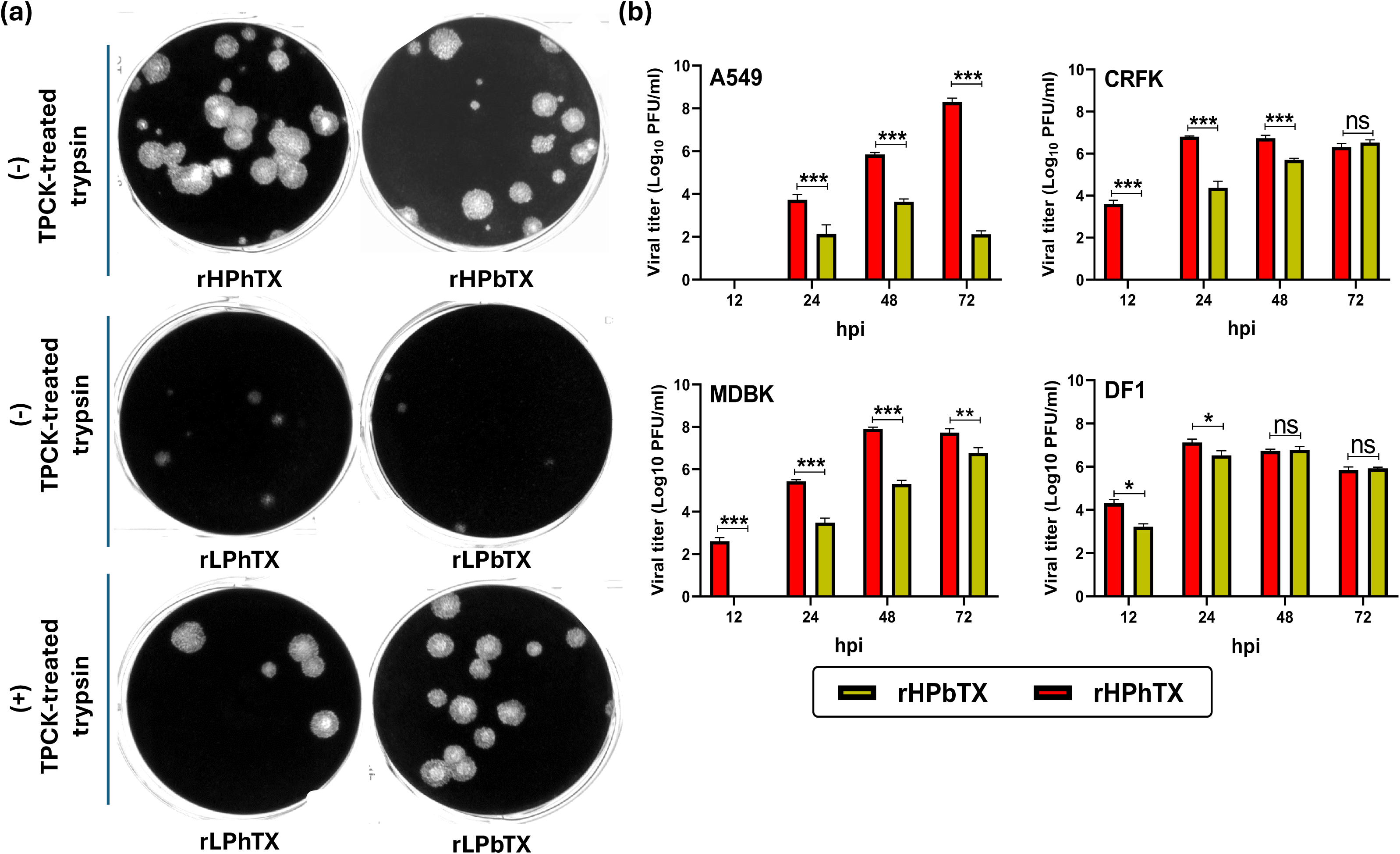
Phenotypic characterization of rHPhTX, rHPbTX, rLPhTX, and rLPbTX. (**a**) Plaque formation in MDCK monolayers with or without TPCK-treated trypsin. (**b**) Growth kinetics of rHPhTX (red) and rHPbTX (green) in A549, MDBK, MDCK, CRFK, and DF-1 cells. Error bars reflect the SD of three independent replicates. Statistical analysis was performed using repeated measures ANOVA, followed by Bonferroni *post hoc* test. The significant differences are indicated (* = *p* < 0.05, ** = *p* < 0.01, *** = *p* < 0.001; non-significant = ns).

Next, we evaluated the ability of currently FDA-approved antivirals to inhibit rHPhTX and rHPbTX. Taking into consideration the identical sequence in the viral M2 protein, we expected that both strains would have comparable susceptibility to M2 amantadine inhibitors. Notably, both strains have no genetic markers that were previously identified to reduce the antiviral activity of the M2 inhibitor amantadine [20]. As expected, the EC_50_ of amantadine was comparable between rHPhTX (0.96 µM) and rHPbTX (0.78 µM) (**Fig 4a**). We next investigated the possible impact of aa variations in PA (K142E, I219L and R497K) and NA (N71S) proteins against PA (baloxavir marboxil) and NA (oseltamivir and zanamivir) inhibitors. Interestingly, baloxavir marboxil (**Fig 4b**), oseltamivir (**Fig 4c**), and zanamivir (**Fig 4d**) exert comparable inhibitory concentrations against rHPhTX and rHPbTX, suggesting that the aa differences in PA or NA do not have an impact on the susceptibility of these viruses to the currently approved FDA antivirals.

**Figure 4.**
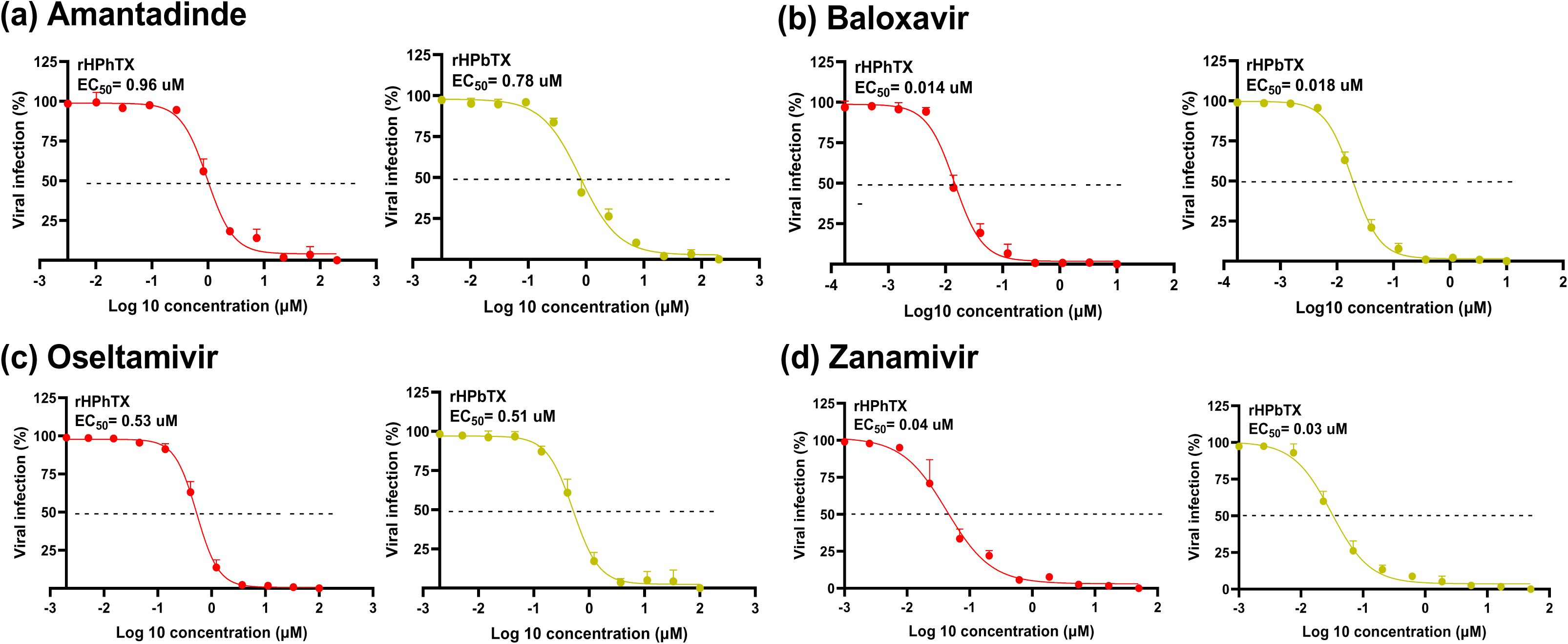
Susceptibility of rHPhTX and rHPbTX to FDA-approved antivirals. MDCK cells were infected with rHPhTX or rHPbTX. After viral absorption, fresh media containing the indicated antivirals was added. Antiviral activities of amantadine (**a**), baloxavir (**b**), oseltamivir (**c**) and zanamivir (**d**) against rHPhTX and rHPbTX were determined at 48 hpi by plotting log(agonist) vs. normalized response—variable slope and the connecting line represents a non-linear regression of the underlying data. Error bars reflect the SD of three independent replicates. The EC_50_ of the indicated antivirals was calculated by plotting log inhibitor versus normalized response (variable slope) and applying the nonlinear regression analyses using GraphPad Prism 9.5.1 software.

### Pathogenicity of rHPhTX, rLPhTX, rHPbTX and rLPbTX in embryonated eggs and C57BL/6J mice

The 50% egg infectious dose (EID_50_) of the HPhTX, rHPhTX, rLPhTX, rHPbTX, and rLPbTX were next determined using 10-day-old embryonated chicken eggs with 10-fold serial dilutions (1-10^4^ PFU) of each recombinant virus. After viral inoculation, embryonated eggs were incubated for 48 h at 37°C and embryo mortality was monitored using egg candler. The egg harvest positivity for influenza virus was detected using standardized hemagglutination assay [21] and the EID_50_ was calculated using the Reed-Muench method [22]. The EID_50_ values for HPhTX and rHPhTX viruses were < 1 PFU, while the rHPbTX was ∼4.64 PFU (**Table 2**). The rLPhTX and rLPbTX viruses have EID_50_ ˃ 10^4^ PFU, in accordance with the expected low pathogenicity of both engineered viruses in avian species.

**Table 2.**
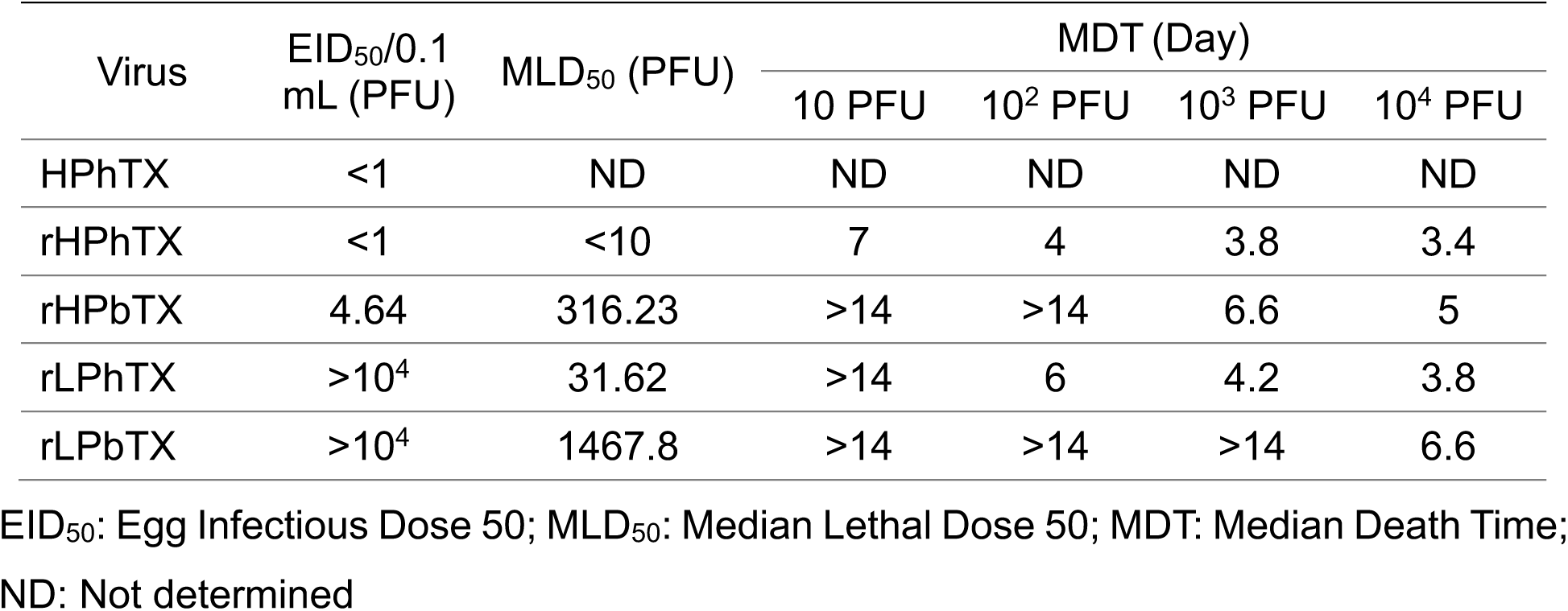
Egg infectious dose 50 (EID_50_), median lethal dose 50 (MLD_50_) and median death time (MDT) of HPhTX, rHPhTX, rLPhTX, rHPbTX and rLPbTX.

We next investigated whether the aa variations between HPhTX and HPbTX can affect the virulence of the highly pathogenic (rHPhTX and rHPbTX) and the low pathogenic (rHPhTX and rHPbTX) viruses in mice. To that end, serial infectious doses (10-10^4^ PFU) of the four recombinant viruses were inoculated into 6-week-old female C57BL/6J mice and animals were monitored for their body weight and survival rates (**Fig 5a**). All mice infected with rHPhTX rapidly lost weight, and all the animals succumbed to viral infection at all applied infectious doses, with a 50% mouse lethal dose (MLD_50_) < 10 PFU and early median death time (MDT) for each infectious dose (**Fig 5a and Table 2**). Mice infected with rLPhTX also showed 100% mortality at high infectious doses (10^2^-10^4^ PFU/mice) with MLD_50_ of ∼31.62 PFU and slightly delayed MDT (Fig 5a and **Table 2**). However, infection with rHPbTX was associated with 100% mortality only for those mice groups infected with high viral titers (10^3^ and 10^4^ PFU), and 20% and 40% mortality at infection doses of 10 and 10^2^ PFU, respectively (**Fig 5b and Table 2**). Unlike the rHPbTX, rLPbTX-infected mice showed no mortality at low infection doses (10 and 10^2^ PFU) and 40% and 100% mortality at 10^3^ and 10^4^ PFU, respectively (**Fig 5b and Table 2**). Compared to rHPhTX (MLD_50_ < 10 PFU) and rLPhTX (MLD_50_ ∼31.62 PFU), both rHPbTX and rLPbTX showed higher MLD_50_ values of ∼316.23 and ∼1,467.8 PFU, respectively.

**Figure 5.**
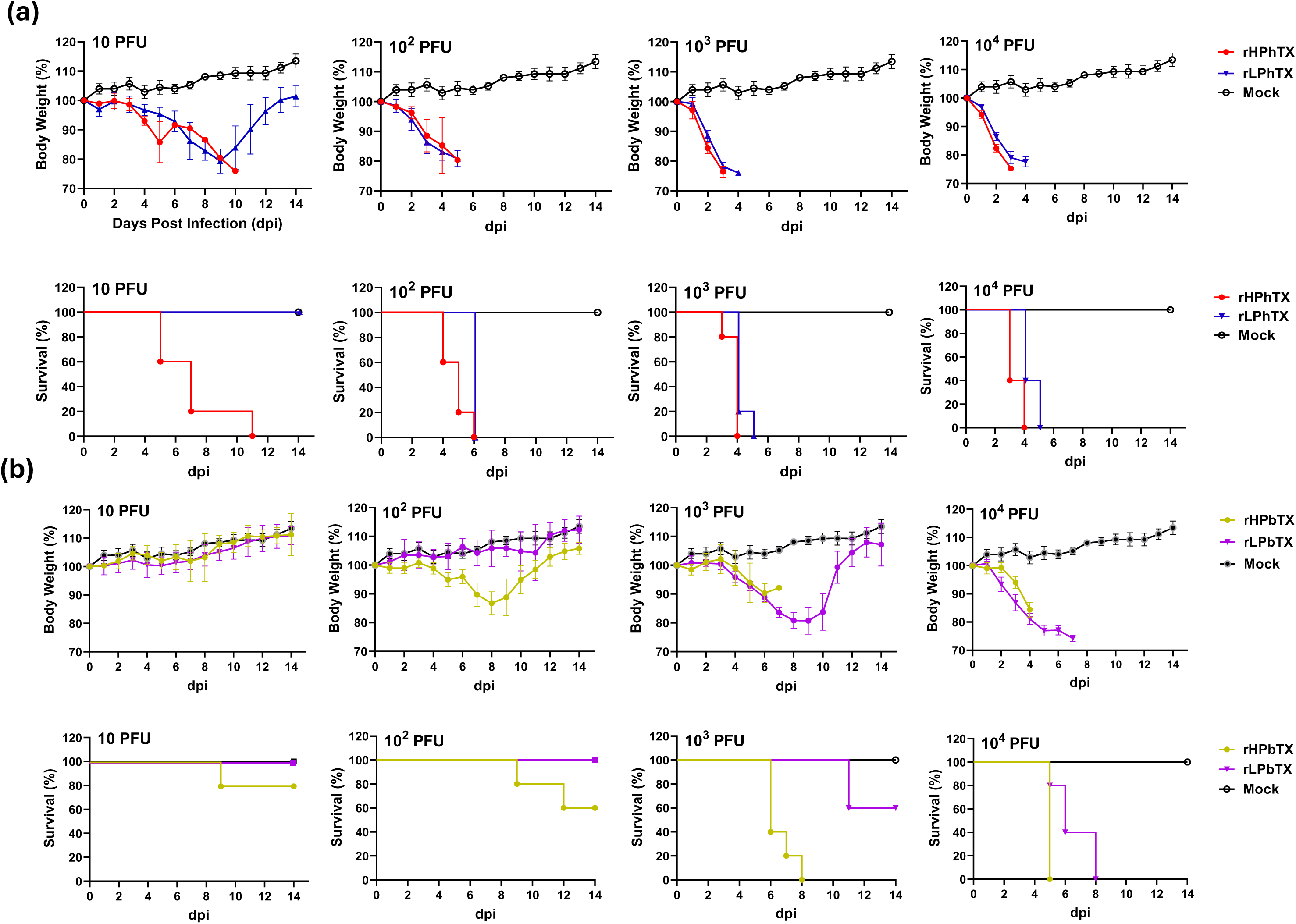
Pathogenicity of rHPhTX, rHPbTX, rLPhTX and rLPbTX in C57BL/6J mice. Percentage (%) of body weight changes (top) and survival curves (bottom) of C57BL/6J mice infected with 10-10^4^ PFU of the rHPhTX and rLPhTX (**a**) or rHPbTX and rLPbTX (b). The mean percent of body weight change (±SEM) is shown. Mice were humanely euthanized when they had lost more than 25% of their initial body weight.

### Viral infection and tissue tropism

Following infection of female C57BL/6J mice with rHPhTX, rLPhTX, rHPbTX, and rLPbTX, two necropsy groups (n=4/group) were euthanized at 2 and 4 dpi to collect nasal turbinate, lung, brain, heart, kidney, spleen, and mammary glands. A group of non-infected mice was used as a control. Half of the collected tissues were individually homogenized and clarified by centrifugation. The virus load in these tissue homogenates was determined by plaque assay. Interestingly, rHPhTX was detected in most tissues of infected mice at 2 and/or 4 dpi (**Fig 6a and Supplementary Fig S1a**). The peak of rHPhTX viral titer was detected in brain tissues at 4 dpi (∼10^5-9^ PFU/mL) according to the viral infection dose. On the same hand, rLPhTX was detected at high titers in nasal turbinate and lung homogenates but had lower titers in brain tissue homogenates at the highest infection doses (10^3^-10^4^ PFU) (**Fig 6a)**, and low to no detection in spleen, kidney, and mammary glands **(Supplementary Fig S1a)**. Similar to the rHPhTX, the rHPbTX could replicate in different organs, including the brain, but the peak of viral titers was lower than those observed in mice infected with rHPhTX (e.g. ∼10^3-6^ PFU/mL) (**Fig 6b and Supplementary Fig S1b**). Unlike the rLPhTX, the rLPbTX could only be detected in the lungs and nasal turbinate from C57BL/6J infected mice (**Fig 6b and Supplementary Fig S1b**). These data demonstrate the contribution of the multibasic cleavage site in the viral HA protein as well as other acquired adaptive substitutions in the polymerases (PB2, PB1, and PA), NA and NS1 proteins in the systemic infection in mice, including tropism to the brain of animals infected with rHPhTX. Notably, and in accordance with the viral replication in the brains, we have been able to observe severe neurological symptoms in mice infected with rHPhTX including paralyses and convulsions, requesting humanized euthanasia.

**Figure 6.**
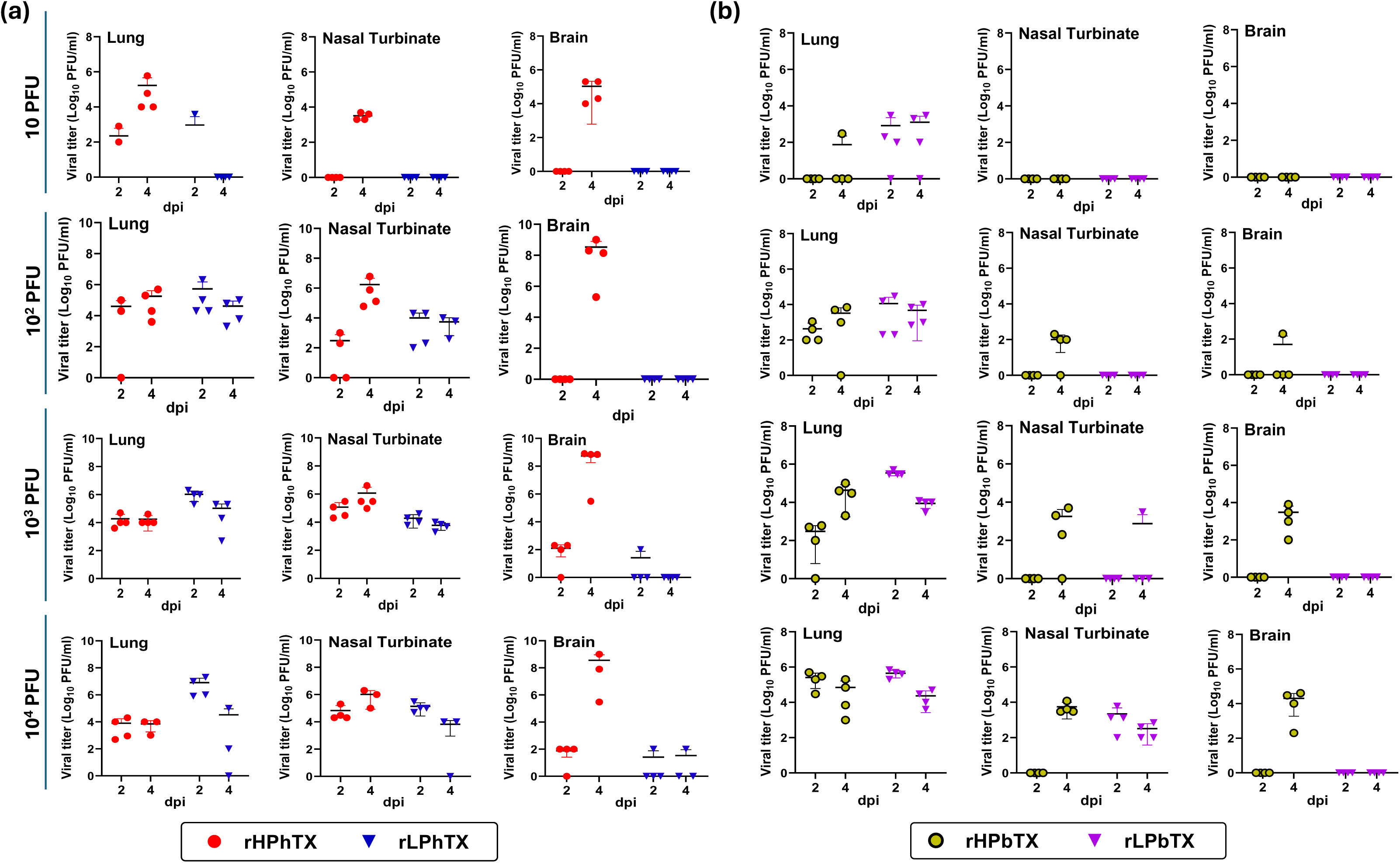
Viral loads in the lung, nasal turbinate, and brain tissues of C57BL/6J mice infected with rHPhTX and rHPbTX. Infectious viral titers in homogenized tissues from mice infected with 10-10^4^ PFU of rHPhTX and rLPhTX (**a**) or rHPbTX and rLPbTX (**b**). The organs were collected at 2 and 4 dpi and were subjected to titration using standard plaque assays.

### Differential histopathology and IHC of infected murine lungs and brains

We next examined the histopathological changes in the lung and brain tissues of C57BL/6J mice infected with rHPhTX and rHPbTX viruses at 4 dpi to confirm that the severe pneumonia progression and neurological complications were associated with efficient viral replication in these organs. The same lung and brain tissue sections were stained for viral antigen by immunohistochemistry (IHC). The histopathological changes attributable to the virus infection, irrespective of the strain of the virus, include varying degrees of interstitial pneumonia characterized by multifocal areas of infiltration of alveolar septa by lymphocytes, plasma cells and lesser macrophages. Small numbers of lymphocytes and macrophages are also found in the alveolar spaces. Occasionally, there is mild fibrin deposition along the septa and some hemorrhage within the areas of inflammation. Blood vessels within the areas of inflammation exhibit endothelial cell hypertrophy and there is mild edema round the larger blood vessels and bronchi. At 4 dpi, the lung tissues from female C57BL/6J mice infected with rHPhTX showed mild inflammatory changes at 10 PFU and multifocal, moderate interstitial pneumonia across 10^2^ PFU to 10^4^ PFU (**Fig 7a**). However, the lung tissue from mice infected with rHPbTX showed mild inflammatory changes at 10 PFU and 10^2^ PFU, and moderate interstitial pneumonia at infection dose of 10^3^ PFU and 10^4^ PFU (**Fig 7a**). Immunohistochemical staining of lungs for viral NP showed positive viral antigen staining in the nucleus and ciliary border of the bronchi and bronchiolar epithelium, strong nuclear staining of the mononuclear cells with the expanded alveolar septa in the areas of inflammation. The extent of viral antigen staining varied significantly between the strains. There was widespread viral antigen with a dose-response pattern varying with the increasing dose of inoculated rHPhTX strain. In contrast, the 10 PFU and 10^2^ PFU groups of rHPbTX showed negligible amount of viral antigen and the 10^3^ PFU and 10^4^ PFU groups of rHPbTX showed dose-dependent increase in viral antigen staining (**Figs 7a and 7b**).

**Figure 7.**
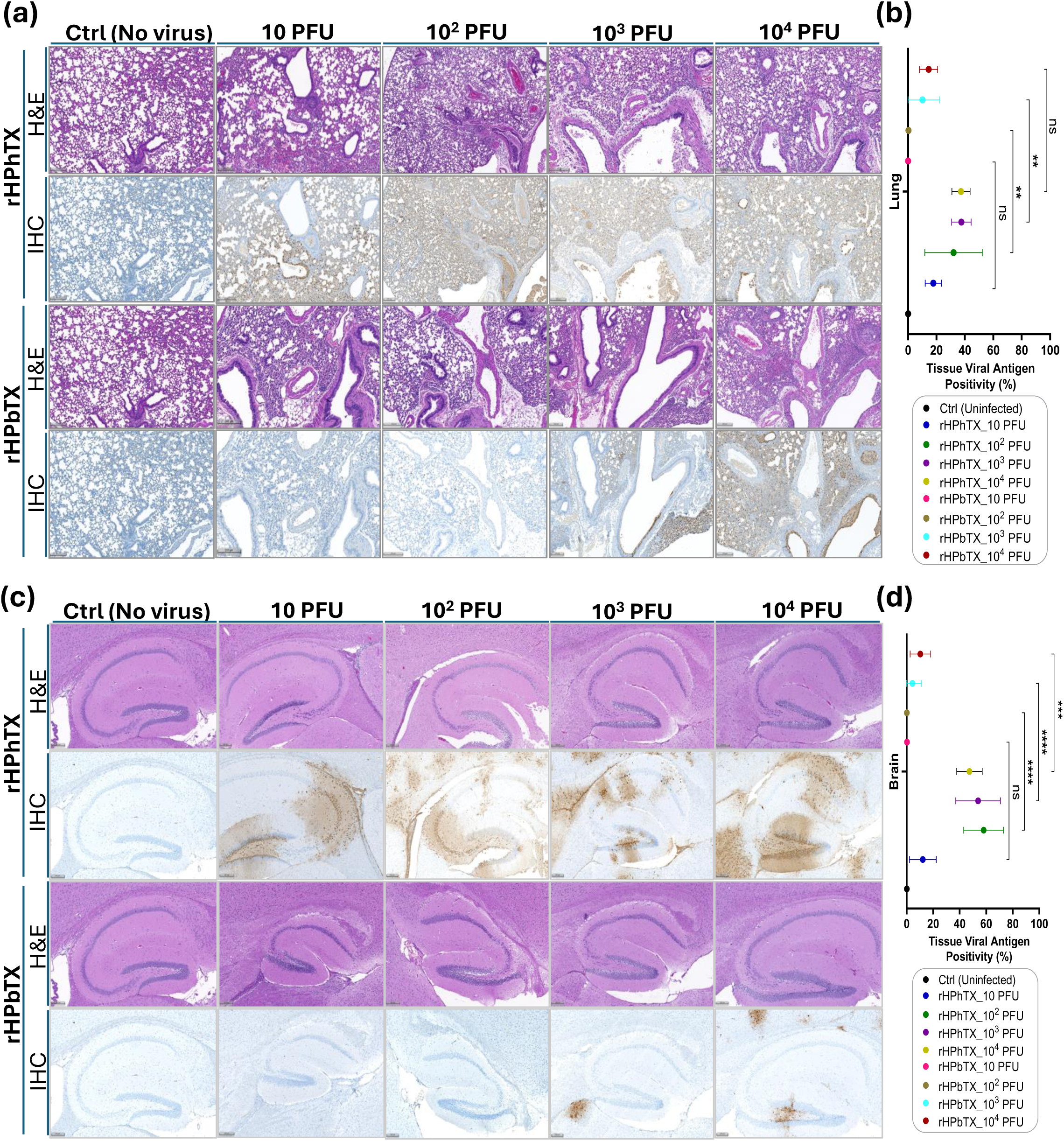
Histopathologic changes and tissue viral NP antigen positivity (%) in the lungs and brains of infected and control C57BL/6J mice. (**a**) Histopathologic changes in murine lung tissues infected with rHPhTX and rHPbTX at different infectious doses (10-10^4^ PFU) against the control uninfected C57BL/6J mice. The upper row corresponding to each virus represents the tissue stained with hematoxylin-eosin (H&E). The bottom row represents the immunohistochemistry (IHC) data of the same part of the tissue (immunostained using an influenza A NP rabbit polyclonal antibody). (**b**) Quantitative assessment of tissue viral antigen positivity (%) in murine lung tissues infected with rHPhTX and rHPbTX at different infectious doses (10-10^4^ PFU) against the control uninfected C57BL/6J mice. (**c**) Histopathologic changes in murine brain tissues infected with rHPhTX and rHPbTX at different infectious doses (10-10^4^ PFU) against the control uninfected C57BL/6J mice. The upper row corresponding to each virus represents the tissue stained with hematoxylin-eosin, while the bottom row represents the immunohistochemistry data of the same part of the tissue. (**d**) Quantitative assessment of tissue viral antigen positivity (%) in murine brain tissues infected with rHPhTX and rHPbTX at different infectious doses (10-10^4^ PFU) against the control uninfected C57BL/6J mice. The pathology overall score was determined following virus quantification in lung and brain tissues of infected C57BL/6J mice (n =4). Data are presented as mean ± SD. Statistical analyses against indicated groups were performed using one-way ANOVA followed by a Tukey post hoc test. The significant differences are indicated (** = *p* < 0.01, *** = *p* < 0.001, **** = *p* < 0.0001; non-significant = ns). Scale bars represent 200μm.

The brain tissues from mice infected with rHPhTX and rHPbTX showed rare to non-significant histopathological changes in all groups observed (**Fig 7c**). The brain tissues were also immunohistochemically stained for viral NP. A strong dose-dependent increase in viral immune straining affecting all parts of the brain was observed in the rHPhTX-infected group. Rare or no viral antigen staining was observed in 10 PFU and 10^2^ PFU groups infected with rHPbTX, and mild antigen staining was observed in 10^3^ and 10^4^ PFU groups infected with rHPbTX. Still, it was significantly less than the comparable dose groups of the rHPhTX-infected mice (**Figs 7c and 7d**). In control groups, the epithelium of bronchi and bronchioles was intact, and the cells stained negative immunohistochemically with no significant pulmonary histopathological changes (**Fig 7**). The full histopathology and IHC from lung and brain tissues from infected C57BL/6J mice are illustrated in **Figs 8a and 8b**, respectively. These data confirm the presence of high viral titers of rHPhTX in infected C57BL/6J mice brain tissue and indicate its broad cellular tropism and neurovirulence, recently observed with similar HPAIV H5N1 strains [23].

**Figure 8.**
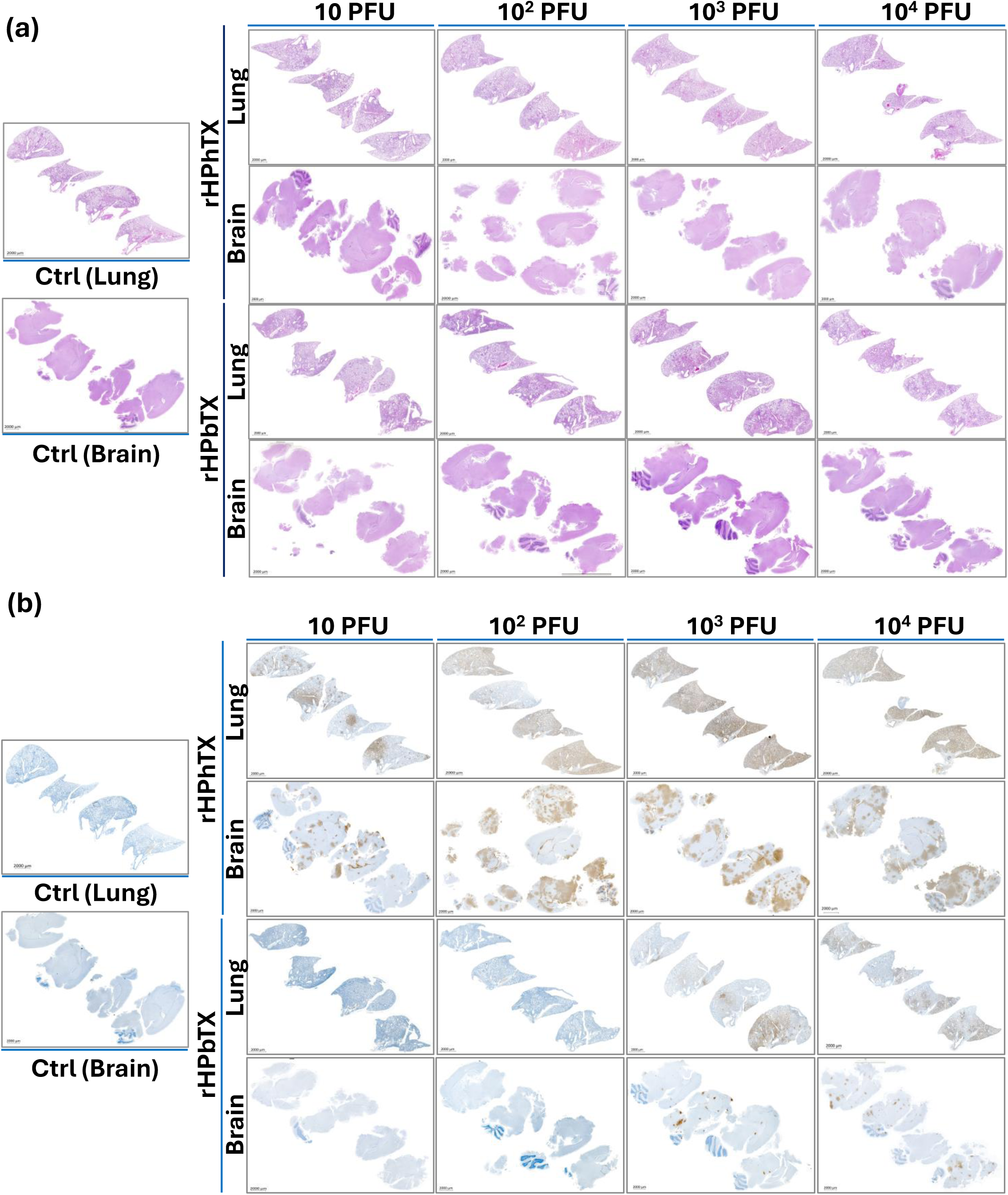
H&E stained and IHC lung and brain tissue sections of control uninfected and rHPhTX- or rHPbTX-infected C57BL/6J mice. (**a**) H&E staining of control uninfected and rHPhTX- or rHPbTX-infected lung and brain tissues of female C57BL/6J mice. (**b**) Influenza NP antigen IHC staining of the control uninfected and rHPhTX- or rHPbTX-infected lung and brain tissues of female C57BL/6J mice. Scale bars represent 2000μm.

### Changes in innate immune responses

Unlike low pathogenic avian influenza viruses (LPAIV), infection with HPAIV H5N1 strains has been associated with the strong induction of a life-threatening cytokine storm (CS). The CS is defined by the increased production of cytokines, chemokines, and interferon-stimulated genes (ISGs) in mammalian hosts including humans, responsible for increased viral pathogenicity [23,24]. To determine whether the differences in morbidity and mortality observed in mice infected with rHPhTX and rHPbTX viruses correlate with cytokine dysregulation, we measured multiple cytokines (IFN-α, IFN-β, IFN-γ, IL-6, IL-17A and TNF-α) (**Fig 9a**) and chemokines (MCP-1 or CXCL2, IP-10 or CXCL-10, and RANTES or CCL5) (**Fig 9b**) in the lung homogenates of C57BL/6J mice infected at 2 and 4 dpi (infection doses of 10, 10^2^, 10^3^, and 10^4^ PFUs). Indeed, all cytokines and chemokines tested were significantly induced by rHPhTX and rHPbTX compared to uninfected C57BL/6J mice either at 2 dpi, 4 dpi, or both time-points. In addition, mice infected with rHPhTX showed significantly higher production of most of the analytes than rHPbTX-infected mice. These differences were more evident at higher infection doses, especially at the earliest time-point (2 dpi). This finding is consistent with the faster and more advanced disease progression and mortality observed in C57BL/6J mice infected with the rHPhTX.

**Figure 9.**
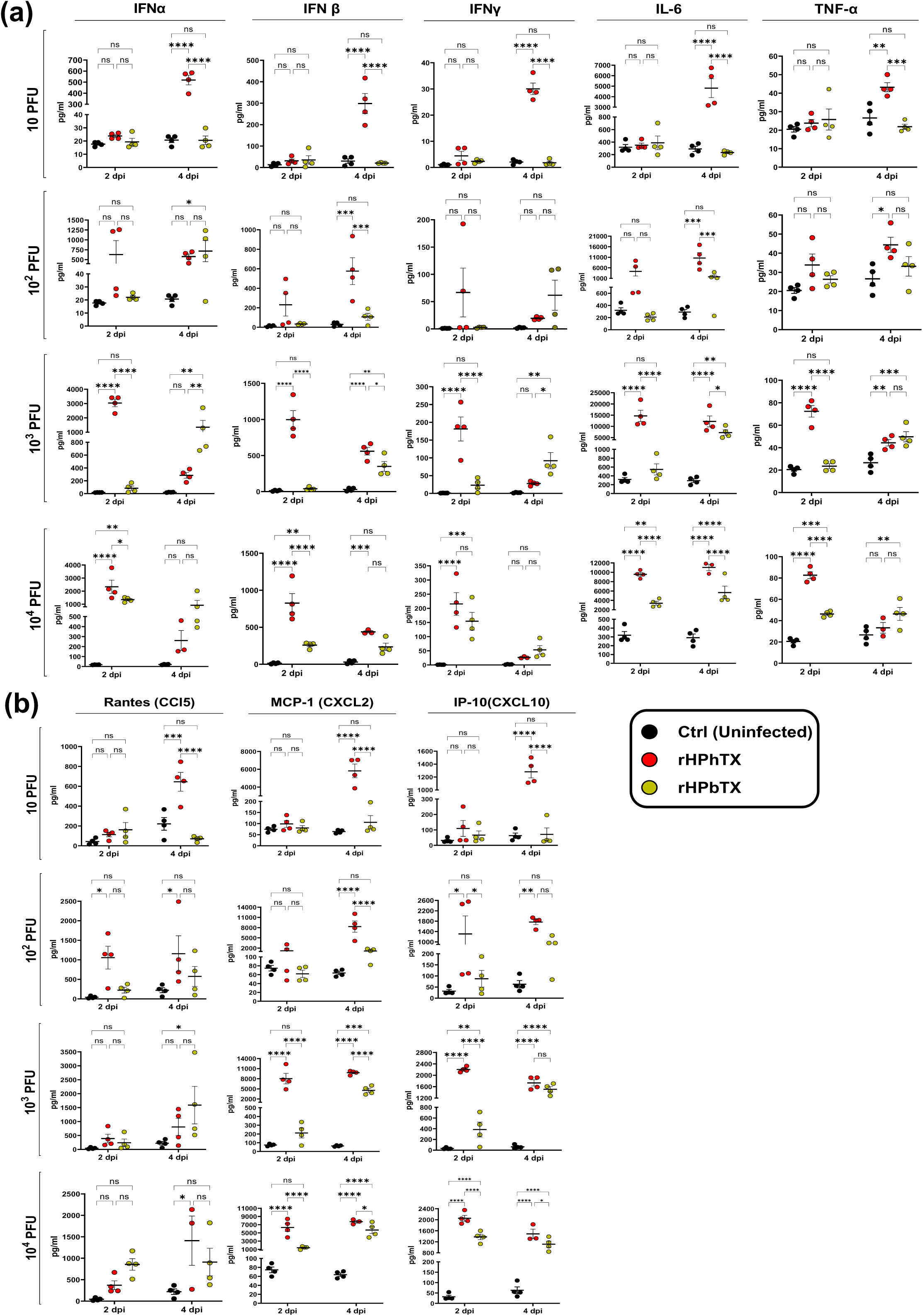
Robust induction of innate immune responses by rHPhTX and rHPbTX in C57BL/6J infected mice. Tissue homogenates from rHPhTX-infected, rHPbTX-infected, or uninfected mice were analyzed at 2 and 4 dpi for secreted cytokines IFN-α, IFN-β, IFN-γ, IL-6, TNF-α, (**a**) and chemokines CCL5, CXCL2 and CXCL-10 (**b**) using an 8-plex bead Luminex assay. Data is shown as M±SEM with n=4 mice/group. A two-way ANOVA with Holm-Sidak’s multiple comparisons test was used to compare groups within each time-point. Statistical significance is reported as *p < 0.05; **p < 0.005; ***p < 0.0005; ****p < 0.00005; ns = non-significant.

## Discussion

Since 2022 until October of 2024, 35 human cases of HPAIV H5N1 clade 2.3.4.4b have been reported in the US, including 34 human cases since April 2024. The 34 human cases were detected following exposure to infected dairy cows (n=19, dairy workers on affected farms), infected birds or poultry (n=14), or no immediately known animal exposure (n=1) [25]. The increasing numbers of human cases since the emergence of the dairy cattle H5N1 outbreaks in March 2024 is alarming, especially with the recent human cases in California from October 3^rd^ to October 10^th^, 2024 [25].

By comparing the first reported cattle-flu-associated human influenza in Texas caused by A/Texas/37/2024(H5N1, HPhTX) to other available isolates from affected dairy cattle herds or humans, we show that the HPhTX has distinct variations in their viral proteins (**Fig 1**). It is not known if HPhTX gained these amino acid changes in cattle, as part of a zoonotic process, or after viral transmission and infection in the human host. The fact that these aa changes are not present in cattle isolates including HPbTX (**Fig 1**), suggest the possibility that the virus rapidly mutates after human infection. In any case, we investigated the impact of these variations on the virus phenotype both *in vitro* and the *in vivo* C57BL/6J mouse model.

To that end, we first generated a recombinant form of HPhTX using reverse genetics and compared its viral replication to the natural HPhTX isolate, showing the same viral kinetics (**Fig 2**). To avoid possible adaptation of the clinical isolate into embryonated eggs and/or MDCK cells used to propagate this virus, the RNA of the natural isolate was subjected to RNAseq and compared to the rHPhTX. Interestingly sequence analyses showed a clonal viral population with only minor nonsynonymous alterations in PA (L655F) and NS1 (P164S) present in the natural isolates (**Supplementary Fig S2**). These variations were not reported in the published sequences of the clinical isolate on GSAID (EPI3171486-93), GenBank (PP577940-47), or the rHPhTX we generated using reverse genetics. Notably, these underrepresented aa changes in PA and NS1 did not change the plaque phenotype in MDCK cell monolayers or viral replication kinetics in different cell lines compared to the rHPhTX generated in the lab (**Fig 2**). To ensure that the recombinant viruses were derived from the rescue plasmids, we introduced a synonymous mutation in PB2 (T2214A →G729G) (**Supplementary Fig S2**) as a genetic tag. In addition, we generated by reverse genetics the cattle-like rHPbTX and the low pathogenic forms of both human and bovine viruses (**Fig 1**).

IAV replication and pathogenicity are polygenic traits that involve multiple viral genes and host factors [3]. The phenotypic comparison of rHPhTX and rHPbTX revealed the advantageous capability of rHPhTX to replicate in different mammalian and avian cell lines, although differences were less pronounced in avian cells (**Fig 3**). HPhTX showed nine aa differences in comparison to the closely related HPbTX virus in PB2 (G362E, E627K and L631M), PB1 (I392V), PA (K142E, I219L and R497K), PA-X (K142E), NA (S71N) and NS1 (Q40R) (**Fig 1**). These genetic changes could have been acquired in the new (human) host due to virus adaptation after infection to improve viral fitness, or in the intermediate animal host (cattle) as part of a zoonotic process to allow cattle-to-human virus transmission, or both [3]. Since all nine substitutions were unique to the HPhTX, they were probably acquired following virus transmission to the farmworker as part of the adaptation process.

The PB2 protein in IAV is one of the crucial viral determinants of virus transmission and adaptation to different mammalian systems [3,17,26]. Unlike the HPhTX, which contains the mammalian adaptive E627K substitution in PB2, the HPbTX contains the mammalian aa substitution M631L that has been frequently reported in the PB2 of dairy cattle H5N1 strains (˃99%). Both substitutions were associated with the increased capability of H5N1 strains to replicate in human cells by enhancing the viral polymerase activity *in vitro* and *in vivo* [17,18,27] allowing the interaction of PB2 with human ANP32A [18,28,29]. Similarly, the K142E substitution in the PA and PA-X proteins of HPhTX was previously reported in human isolates of A/Hong Kong/156/97(H5N1) and A/Vietnam/1203/2004(H5N1) and it has been found to be associated with high polymerase activity in mammalian cells [30], suggesting a role for this PA residue in virus adaptation to humans. The potential role of the other aa substitutions in the PB2 (G362E), PB1 (I392V) and PA (I219L and R497K) have not been described.

The nonstructural protein 1 (NS1) of IAV has an N-terminal RNA-binding domain (RBD) that binds to double stranded viral RNA (dsRNA) and sequester cellular mRNAs species to antagonize the host’s antiviral response [31]. The aa residues R38 and K41 are essential for binding viral dsRNA and cellular mRNAs. Of interest is the Q40R substitution in the NS1 of HPhTX that was not observed in other B3.13 sequences from dairy cattle or human H5N1 strains (**Fig 1**). This mutation resides in the RNA-binding domain of the NS1 between the essential R38 and K41 aa, required to sit and stabilize the interaction with the dsRNAs [32], and its contribution to the ability of NS1 to control host cell shutoff and/or antagonize innate immunity remains to be investigated.

Unlike the NA of the HPhTX that contains S71N, the main cattle outbreak clade, including the cattle-flu-related human cases, have NA proteins with N71S [33] (**Fig 1**). NA N71S substitution has been reported in immunocompromised patients with seasonal influenza infection in combination with the known E119V resistance marker after long-term treatment (139 days) with cumulative zanamivir therapy (E119V (100%) and N71S (100%)) [34]. It is unclear whether this substitution emerged as an Oseltamivir resistance marker or acquired within a long adaptation process in an immunocompromised mammalian system. To assess the possible contribution of aa differences in the human isolate to antiviral resistance, we tested the FDA-approved endonuclease (baloxavir marboxil), NA (oseltamivir and zanamivir) and M2 (Amantadine) inhibitors against rHPhTX and rHPbTX (**Fig 4**). As expected, the susceptibility to amantadine was comparable between both viruses since they share the same sequence in the M2 protein, and they do not present any aa changes previously identified to confer resistance to amantadine/rimantadine. Both viruses were also inhibited by the PA (baloxavir) and NA (oseltamivir and zanamivir) inhibitors. Comparable EC_50_ values of these FDA-approved drugs against rHPhTX and rHPbTX indicate that the aa substitutions in PA and NA are not contributing to resistance episodes to available drugs against influenza (**Fig 4**).

The virulence of dairy cattle H5N1 strains was recently investigated in female BALB/cJ mice [8], and it was reported that all mice infected with 10^3^ PFU, or higher, of the A/dairy cattle/New Mexico/A240920343-93/2024(H5N1) succumbed to viral infection, with some mice infected with 10 and 10^2^ PFU surviving infection. These data were consistent with our findings, where all rHPbTX-infected C57BL/6J mice succumbed to infection using 10^3^-10^4^ PFU, while 80% and 60% of the infected mice with 10 and 10^2^ PFU survived infection until 14 dpi (**Fig 5**). Taking into consideration that the used bovine H5N1 isolates and mice strains are different, both studies showed different MLD_50_ values for the bovine H5N1 strains [8]. Interestingly, the rHPhTX led to higher virulence in C57BL/6J mice, where all mice succumbed to the infection at infectious dose ≥ 10 PFU (**Fig 5**) with a significant increased productions of inflammatory cytokines and chemokines in the lungs of infected C57BL/6J mice (**Fig 9**). Conversely, both low pathogenic forms, rLPhTX and rLPbTX, led to slightly delayed mortality compared to the rHPhTX and rHPbTX counterparts (**Fig 5**). This suggests that HA multibasic cleavage site in the human and bovine strains contributes to the increased viral replication and pathogenicity in C57BL/6J mice. These results further confirm the role of this site, cleaved by furin-like proteases when multibasic, in the pathogenicity of H5N1 viruses. Besides, it is apparent that the increased virulence rLPhTX, compared to rLPbTX, is because the nine aa substitutions that are enough to increase the ability of rLPhTX to exploit the mammalian host, even in the absence of the HA multibasic cleavage site.

Unlike human and other avian influenza viruses, a unique feature of infections with HPAI H5N1 strains is that they are associated with severe neurological complications in avian and mammalian hosts. In the current study, mice infected with rHPhTX showed strong neurological symptoms shortly before death or euthanasia. These symptoms were dose-dependent and associated with higher viral loads in the brain tissues of mice infected with rHPhTX as determined by viral titrations (**Fig 6)** and IHC detection of viral antigens (**Figs 7 and 8**). Intriguingly, despite high pathogenicity in mice with neurological consequences, all documented human infections with bovine H5N1 viruses did result in mainly conjunctivitis with non-severe symptoms. At this moment, it is unclear if this is due to differences between mice and humans with respect to the pathogenesis of H5N1 viruses, or to the acquisition of the virus infection through the eye in all the human reported cases.

Collectively, our study demonstrates that the rHPhTX with acquired adaptative substitutions that improved replication efficiency and ability to induce proinflammatory innate immunity and pathogenicity in C57BL/6J mice. These adaptations might increase the potential of H5N1 viruses to acquire the ability to transmit among humans. As a result, human H5N1 strains must be closely monitored and assessed for their public health risk, and efforts should be made to eradicate the H5N1 from cows, in order to avoid further human H5N1 infections by these already mammalian adapted H5N1 viruses.

## Acknowledgments

We thank the Histology unit team staff, Renee Escalona and Colin Chuba, for their assistance in tissue staining/immunostaining experiments, and the Cell Biology Core Lab at Texas Biomedical Research Institute for assistance with the multiplex cytokine assay. We are grateful to Todd Davis and Han Di at the Virology, Surveillance and Diagnosis Branch, Influenza Division, The Centers for Disease Control and Prevention (CDC) for providing the natural isolate of influenza A/Texas/37/2024(H5N1). We also thank Daniel Perez at the Animal Health Research Center, Center for Vaccines and Immunology, Department of Population Health, Poultry Diagnostic and Research Center for providing MDBK cells.

## Authors contributions

Conceptualization: A.M. and L.M-S.; Methodology: A.M., R.S.B., A.A-G., R.A.E., V.S., E.M.C., Y.A., H.R., N.J., M.B., C.Y., and A.C.; Data collection and interpretation: A.M., R.S.B., A.A-G., V.S., A.G-S., J.B.T., and L.M-S.; Funding acquisition and resources: A.G-S., J.B.T., and L.M-S.; Writing—original draft preparation: A.M. and L.M-S.; Writing— review and editing: all authors have read and agreed to the published version of the manuscript.

## Figure legends

**Supplementary Fig S1. Viral loads in heart, spleen, kidney, and mammary glands tissues of C57BL/6J mice.** Infectious viral titers in homogenized tissues from mice infected with 10-10^4^ PFU of rHPhTX and rLPhTX (**a**) or rHPbTX and rLPbTX (**b**). The organs (n=4) were collected at 2 and 4 dpi and were subjected to titration using plaque infectivity assay.

**Supplementary Fig S2.** (**a**) HPhTX genome coverage by next-generation sequencing (NGS). (**b**) Clonal populations of the virus with minor nonsynonymous alterations in PA (L655F) and NS1 (P164S) were detected in the adapted natural isolate not previously reported in the published sequences (GenBank accession numbers of PP577940-47). The nonsynonymous mutation T2214A was introduced in the rHPhTX PB2 segment as a genetic tag to differentiate from the natural HPhTX isolate.

