## Supplementary figures and images for "Replication Kinetics, Pathogenicity and Virus-induced Cellular Responses of Cattle-origin Influenza A(H5N1) Isolates from Texas, United States"

### Supplemental Figure 1

#### Supplementary Figure S1

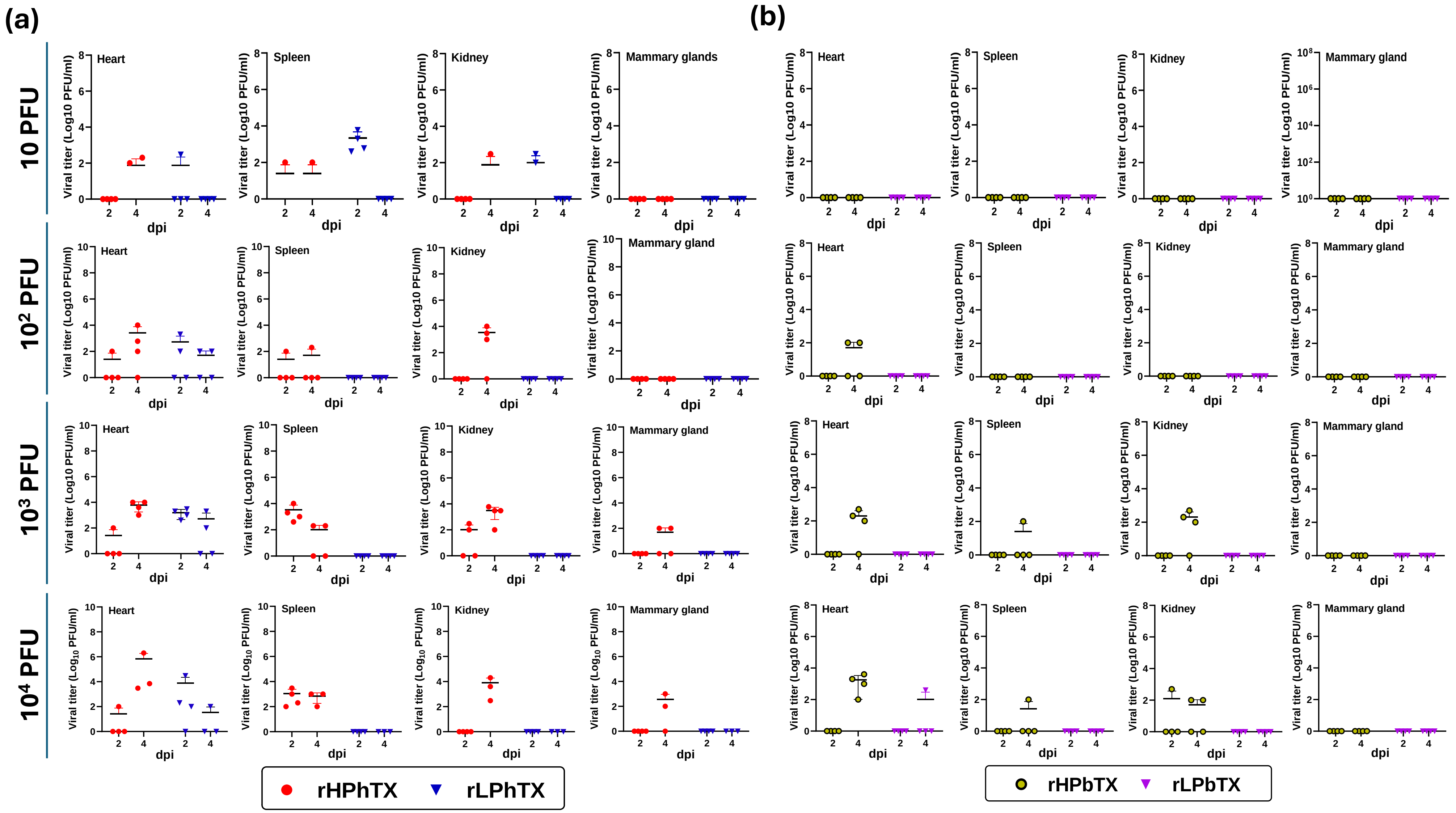

### Supplemental Figure 2

**Supplementary Figure S2**

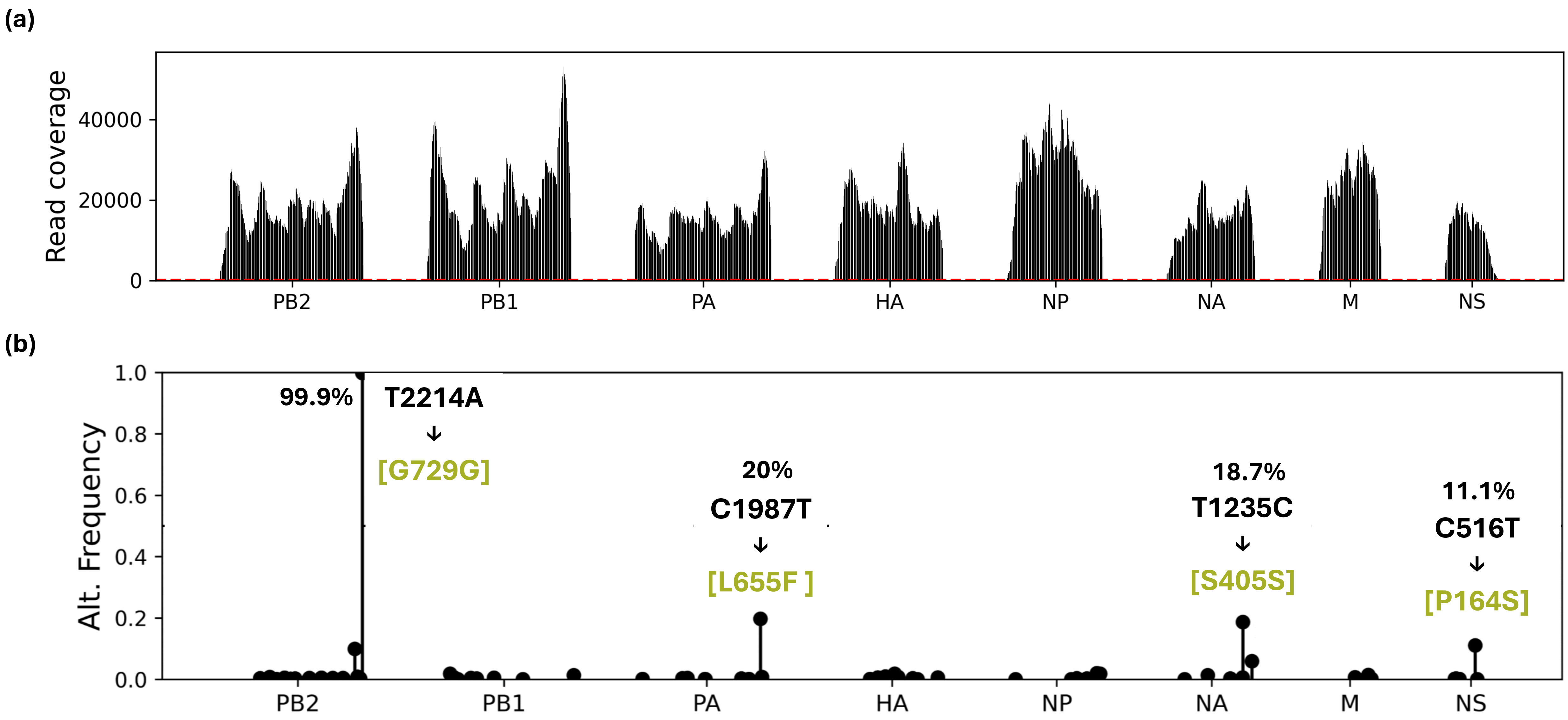
